# Photoreceptor inputs to pupil control

**DOI:** 10.1101/624759

**Authors:** Manuel Spitschan

## Abstract

The size of the pupil depends on light level. Watson & Yellott (2012) developed a unified formula to predict pupil size from luminance, field diameter, age, and number of eyes. Luminance reflects input from the L and M cones in the retina but ignores the contribution of intrinsically photosensitive retinal ganglion cells (ipRGCs) expressing the photopigment melanopsin, which are known to control the size of the pupil. We discuss the role of melanopsin in controlling pupil size by reanalysing an extant data set. We confirm that melanopsin-weighted quantities, in conjunction with Watson & Yellott’s formula, adequately model intensity-dependent pupil size. We discuss the contributions of other photoreceptors into pupil control.

---

In a paper adequately described as a *tour de force*, Watson and Yellott (2012) developed a unified formula to predict pupil size from luminance, field diameter, age, and number of eyes. This letter concerns the parametrisation of the retinal intensity, which in Watson and Yellott’s model is given in terms of luminance, i.e. the radiance of the stimulus weighted by the photopic luminosity curve V(λ). V(λ) corresponds to a mixture of the L and M cones in the retina, thereby largely ignoring the potential role of S cones, rods, and the intrinsically photosensitive retinal ganglion cells (ipRGCs) expressing the photopigment melanopsin (Dacey et al., 2005; Gamlin et al., 2007; Provencio et al., 2000).

The observation that V(λ)-weighted quantities do not predict pupil size is not new (Berman, Jewett, Fein, Saika, & Ashford, 1990; Krastel, Alexandridis, & Gertz, 1985). Already in 1962, Bouma (1962) noted that the spectral sensitivity of pupil control is neither V(λ) nor the rod-based V’(λ), interjecting that the outcome of his experiments “may turn out to be related to other adaptive processes in the human eye”. Bouma himself modelled the spectral sensitivity as a combination of S cones and rods. We know now that steady-state pupil size is largely controlled by melanopsin.

To test if Bouma’s data is consistent with melanopsin-based pupil control, we reanalysed the intensity-response curves from Bouma (1962) as follows. We first extracted the data from Bouma’s Figure 1, as shown in our Figures **1*A*** and **1*B*** using WebPlotDigitizer, https://automeris.io/WebPlotDigitizer/). For monochromatic lights, which we assumed Bouma used, it is simple to convert the reported V(λ)-weighted luminous flux into a melanopsin-weighted radiant flux (CIE, 2018). As radiant flux describes the total amount of energy emitted by a source, it is not an appropriate measure to describe corneal or retinal intensity, so the absolute quantities are not informative unless a geometry is specified. Allowing for fixed but arbitrary horizontal shift, the data for all wavelengths now coincide, except for long-wavelength lights (Fig. **1*C***). In addition, Watson and Yellott’s formula (red line) accounts well for the shape of the pupil response as a function of melanopic radiant flux.

**Figure 1.**
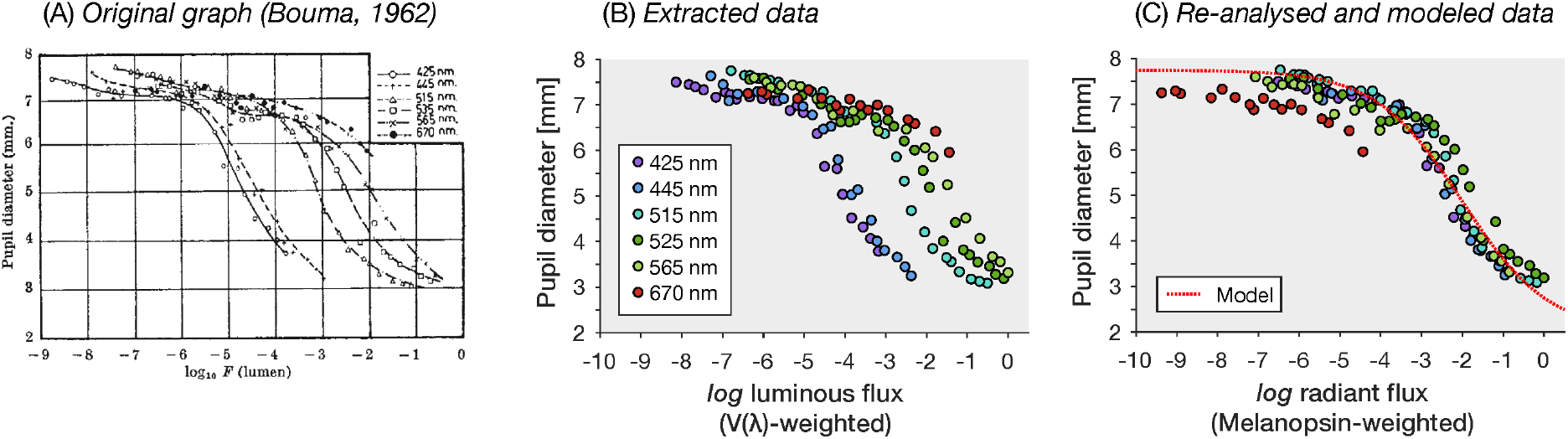
***A*** Original graph from Bouma (1962) relating luminous flux to pupil diameter in millimeters. ***B*** Replotted extracted pupil size data. ***C*** Data replotted in terms of normalized melanopic radiant flux, along with the unified formula by Watson and Yellott (2012), allowing for a horizontal shift aligning the data with the curve.

One notable and systematic deviation occurs for the 670 nm data points, which a melanopsin-exclusive model appears not to predict well. This suggests that melanopsin is not the only photoreceptor controlling steady-state pupil size. This is indeed not surprising, as melanopsin-containing retinal ganglion cells receive cone and rod inputs (Dacey et al., 2005).

Indeed, there is now a good body of evidence that all photoreceptors can control the diameter of the pupil. The best evidence comes from studies examining pupil size using the method of silent substitution, in which pairs of lights are alternated such that only one photoreceptor class is stimulated (Estévez & Spekreijse, 1982; Spitschan & Woelders, 2018). Studies examining pupil control using this method are given in **Table 1**.

**Table 1:** Evidence of photoreceptor contributions to pupil control.

| Photoreceptor class | Reference |
| --- | --- |
| Melanopsin | Tsujimura, Ukai, Ohama, Nuruki, and Yunokuchi (2010)<br>Vienot, Bailacq, and Rohellec (2010)<br>Tsujimura and Tokuda (2011)<br>Spitschan et al. (2014)<br>Cao, Nicandro, and Barrionuevo (2015)<br>Barrionuevo and Cao (2016)<br>Spitschan et al. (2017)<br>Zele, Feigl, Adhikari, Maynard, and Cao (2018) |
| L cone | Spitschan et al. (2014) (L+M)<br>Spitschan et al. (2017) (L+M+S)<br>Barrionuevo and Cao (2016)<br>Murray, Kremers, McKeefry, and Parry (2018)<br>Woelders et al. (2018) |
| M cone | Spitschan et al. (2014) (L+M)<br>Spitschan et al. (2017) (L+M+S)<br>Barrionuevo and Cao (2016)<br>Murray et al. (2018)<br>Woelders et al. (2018) |
| S cone | Spitschan et al. (2014)<br>Spitschan et al. (2017)<br>Barrionuevo and Cao (2016)<br>Cao et al. (2015)<br>Murray et al. (2018)<br>Woelders et al. (2018) |
| Rods | Barrionuevo, McAnany, Zele, and Cao (2018)<br>Barrionuevo et al. (2014) |

A key realisation is that while all photoreceptors may contribute to controlling the pupil size, the when and how is important. For example, due to rod saturation (Aguilar & Stiles, 1954), rods are not expected to contribute to pupil control at photopic light levels. The temporal regimes in which the photoreceptors contribute are also different. Notably, L+M stimulation is band-pass, while S cones and melanopsin are tuned to low frequencies in driving the pupil (Spitschan, Jain, Brainard, & Aguirre, 2014). McDougal and Gamlin (2010) found that cones and rods account for pupil constriction between 1 and 10 seconds from the onset of the light exposure, at 100 seconds, pupil size is largely controlled by melanopsin with some contribution from the rods.

To what extent does Watson and Yellott’s use of luminance as an input parameter call into question the generalizability of their model? From first principles, differences between V(λ)-weighted and melanopic quantities are largest with monochromatic lights. But we typically do not live under monochromatic illumination. We explored this question by examining the range of melanopic irradiances at a fixed illuminance. In other words, how wrong would we be if we continued using V(λ)-weighted quantities to predict pupil size? Using a database of 401 polychromatic (“white”) illuminant spectra (Houser, Wei, David, Krames, & Shen, 2013), we calculated the range of melanopic irradiance while keeping the photopic illuminance fixed at 100 lux (Figure **2*A***). Across all 401 spectra, a 100 lux light source has a melanopic irradiance of 75.5±23.4 mW/m^2^. Crucially, the melanopic irradiance of an illuminant at 100 lux depends also on the correlated colour temperature (CCT) of the source (Fig. **2*B***), with higher-CCT illuminants generally having a higher melanopic irradiance. Irrespective of CCT, the range of melanopic irradiances is between 20.4 and 164 melanopic mW/m^2^, i.e. in the worst case a factor of ~8. In other words, using a V(λ)-based pupil formula could lead to a misrepresentation of the retinal intensity by up to one order of magnitude (log_10_(8) = 0.9), which manifests in a horizontal shift of the intensity response curve in Fig. **1*A-C***.

**Figure 2.**
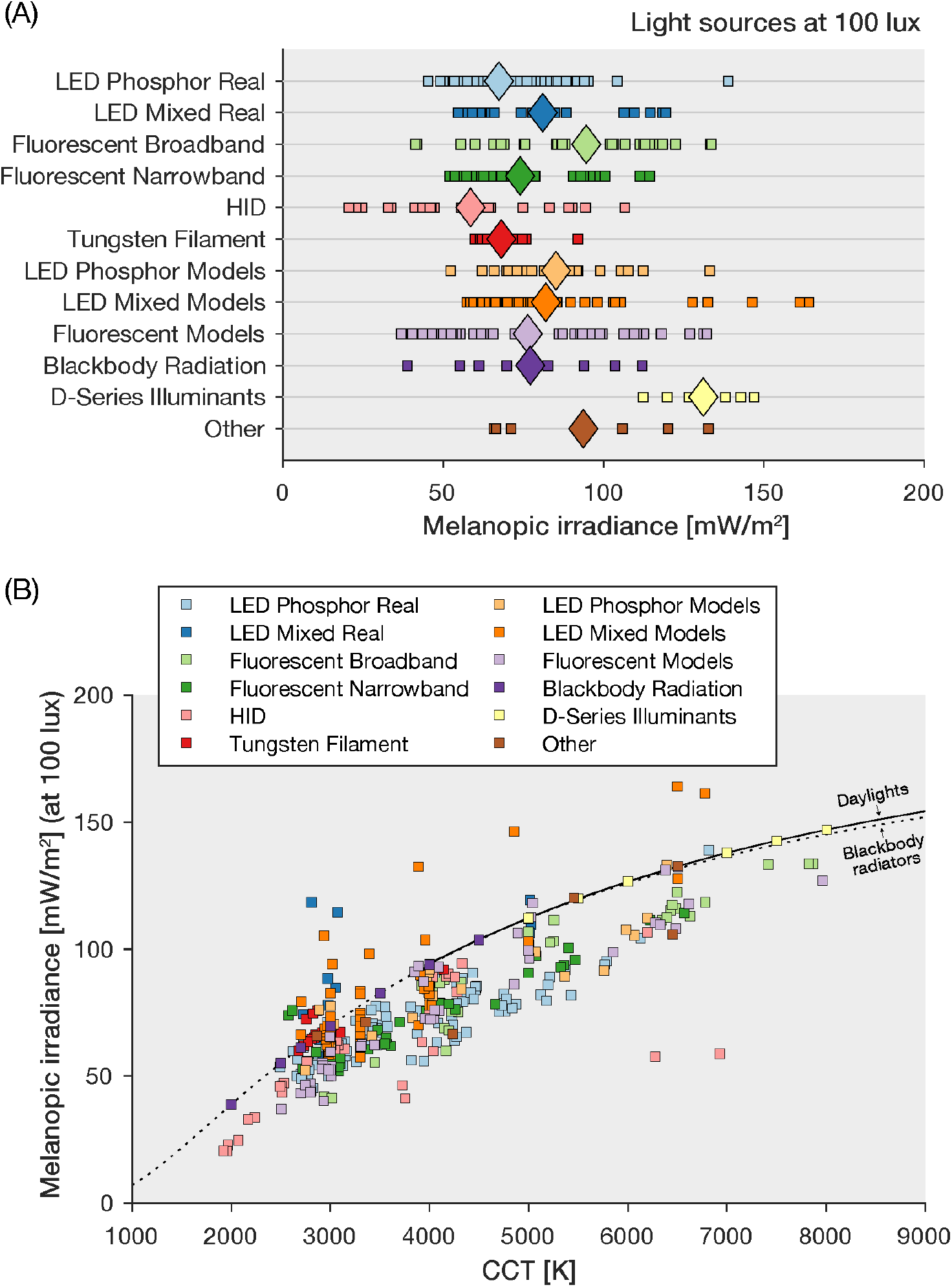
***A*** Variability of the melanopic irradiance of 401 polychromatic “white” light sources (Houser et al., 2013) at 100 lux. Individual sources are represented as individual dots. Large diamonds are the per-category mean. Houser et al. (2013) considered both real and theoretical (i.e. model-based) light sources. Details on the nomenclature and provenance of the light source can be found in (Houser et al., 2013) ***B*** Melanopic irradiance as a function of the correlated colour temperature (CCT). Continuous line indicates daylights; dashed line indicates blackbody radiators.

The degree of misestimation of pupil size from a V(λ)-based model depends on the retinal intensity (as the curve is non-linear). It is also conceivable that the diversity in pupil formulæ found by Watson and Yellott (2012) could simply reflect the fact that previous investigators used different spectral power distributions, which had the same (il)luminance but differed in their melanopic (ir)radiance.

Whether or not the worst-case misprediction by using a V(λ)-weighted quantity has tangible consequences depends on the application. Predicting pupil size in a psychophysical experiment at mesopic light levels requires less stringent estimation of retinal intensity than safety-critical calculations.

A recent study reported an attempt to derive a formula for predicting pupil size from melanopsin activation but only focused on a rather narrow luminance range (50-300 cd/m^2^) (Rao, Chan, & Zhu, 2017). While this is a good start, it might be a useful empirical exercise to collect natural pupil sizes under a large range of illumination conditions (indoors, outdoors) under natural behaviour with conjoint spectral measurements. Our analysis of pupil size as a function of melanopic retinal intensity provides a starting point for predicting pupil size from the spectral properties of a scene.

## Code and data

All data and code, including a MATLAB implementation of Watson & Yellott’s formula, are available at https://github.com/spitschan/Spitschan2019_JOV.

